# Cargo Transport Shapes the Spatial Organization of a Microbial Community

**DOI:** 10.1101/318600

**Authors:** Abhishek Shrivastava, Visha K. Patel, Yisha Tang, Susan Connolly Yost, Floyd E. Dewhirst, Howard C. Berg

**Author notes:** Author Contributions: A.S., F.E.D., and H.C.B. designed the experiments. A.S., V.K.P, Y.T., and S.C.Y. performed the experiments. A.S. and H.C.B. wrote the paper. This article is a PNAS contribution.

## Abstract

The human microbiome is an assemblage of diverse bacteria that interact with one another to form communities. Bacteria in a given community are arranged in a three-dimensional matrix with many degrees of freedom. Snapshots of the community display well-defined structures, but the steps required for their assembly are not understood. Here, we show that this construction is carried out with the help of gliding bacteria. Gliding is defined as the motion of cells over a solid or semi-solid surface without the necessity of growth or the aid of pili or flagella. Genomic analysis suggests that gliding bacteria are present in human microbial communities. We focus on *Capnocytophaga gingivalis* which is present in abundance in the human oral microbiome. Tracking of fluorescently-labeled single cells and of gas bubbles carried by fluid flow shows that swarms of *C. gingivalis* are layered, with cells in the upper layers moving more rapidly than those in the lower layers. Thus, cells also glide on top of one another. Cells of non-motile bacterial species attach to the surface of *C. gingivalis* and are propelled as cargo. The cargo cell moves along the length of a *C. gingivalis* cell, looping from one pole to the other. Multi-color fluorescent spectral imaging of cells of different live but non-motile bacterial species reveals their long-range transport in a polymicrobial community. A swarm of *C. gingivalis* transports some non-motile bacterial species more efficiently than others and helps shape the spatial organization of a polymicrobial community.

**Significance:** We describe a situation in which bacteria typical of the human oral microbiome are organized spatially by gliding cells, species of *Capnocytophaga*, that move backwards and forwards over the substratum. The mobile adhesins that pull the cells over the substratum also attach to cells of non-motile bacterial species, which are carried up and down the motile cells as cargo. The synchronized transport of non-motile cargo bacteria helps shape a polymicrobial community.

## Introduction

Bacteria that attach to and colonize a surface form some of the most stable species of the human microbiome. Rapid availability of nutrition, protection from antibiotics, and long-term colonization are advantages of attaining a specific spatial niche within a microbial community. The mechanisms that drive interactions and guide the architecture of microbial communities are unclear. Active motion via the help of a molecular motor and guidance by a chemotaxis pathway enables some bacteria to sense their environment and find a spatial niche (1, 2). While the chemotaxis ability of a bacterium improves surface colonization (3, 4), motile bacteria do not always dominate a microbiome. For example, out of the thirteen abundant genus of bacteria that colonize the human gingival region (5), bacteria of three genus *Capnocyophaga*: gliding motility (6), *Neisseria*: twitching motility (7), and *Lautropia*: flagellar motility (8) are known to be motile. Non-motile bacteria can reach a spatial niche via diffusion or fluid flow. However, diffusion is not an efficient means of transport for an object of micron size, and fluid flows can be erratic: a sphere of radius 1 μm suspended in water at room temperature takes ~2.5 hours to diffuse a distance of 60 µm (9). Another way by which non-motile bacteria can be transported is by adhering to motile microorganisms. Only a few examples of this process are known (10–12), and its overall significance is unclear. Here, we describe a mechanism in which motile but non-flagellated bacteria of the human oral microbiome carry non-motile bacteria as cargo.

Collective motion is observed in biological objects of all sizes, such as bacteria, insects, birds, and fish (13, 14). The transition of an isolated cell to a swarm can be broadly explained by models based on phase transitions (14). Variations in adhesion forces, the mode of motility, and a bias in the direction of motion govern behavior of a particular species. Swarms of flagellated bacteria comprise dense groups of cells that swim in a thin layer of liquid close to a surface, without adhering to that surface (15, 16). Bacteria with type 4 pili are propelled as the pili attach, retract, and detach from the surface, performing twitching motility. Bacteria that use mobile cell-surface adhesins are called gliding bacteria. Bacterial gliding occurs when bacteria move in close contact with an external surface, a process that requires continuous expenditure of energy (17–19).

Bacteria of the phyla Bacteroidetes are abundant in human gut and oral microbiomes, and many members of this phyla have the ability to navigate surfaces via gliding motility. The Bacteroidetes represent about half of the bacterial population of the human gut microbiome (20,21) and changes in their concentration correlates with diseases such as obesity, diabetes and colitis (22–24). The Type IX Protein Secretion System (T9SS) is found in many Bacteroidetes and is required for the secretion of proteins such as cell-surface adhesins, chitinase, cellulase, and proteases. T9SS is found in both motile and non-motile Bacteroidetes. The core T9SS proteins are SprA, SprE, SprT, and GldK-N. Bacteria with a cell-surface adhesin SprB, motility proteins GldA-B, GldD, GldF-J, and a functional T9SS are able to move over external surfaces (25).

The human oral microbiome is a catalog of microorganisms from lips, teeth, gingiva, tongue, cheek, palate, and contiguous extensions of the oral cavity (26). Bacteroidetes of the genus *Porphyromonas, Prevotella*, and *Capnocytophaga* are found in abundance in the human oral microbiome, and they contain the genes to encode the T9SS. Several reports show the abundance of *Capnocytophaga* sp. in different sites of the human oral microbiome (5, 27–29). Some *Capnocytophaga* sp. isolated from the human oral microbiome form spreading colonies (6), and cells lacking the T9SS are deficient in gliding (30). *Capnocytophaga sp.* isolated from the human oral microbiome possess the genes to encode the T9SS and its associated gliding machinery. This suggests that the mechanism of gliding found in the phyla Bacteroidetes is prevalent in the genus *Capnocytophaga*. However, no experimental data exist about the mechanism for the motion of single cells of the genus *Capnocytophaga*, nor is anything known about their collective behavior. Recently, via fluorescent labeling of fixed samples of human oral biofilms, Welch *et al.* showed the presence of a structured and multigenus consortia within human oral microbial communities that contained bacteria of the genus *Capnocytophaga* (5). How a non-ordered community reaches an ordered state and arranges itself into spatial structures is not clear. Sequence and imaging data showed that bacteria of the *Capnocytophaga* genus are found in abundance in both human supragingival and subgingival biofilms. Why the members of a genus that represents gliding bacteria are stable and abundant in the oral microbiome of healthy humans is not known. Via spectral imaging of live cells, we study the development of a polymicrobial community that contains eight abundant bacterial species of a human subgingival biofilm. We show that swarming *Capnocytophaga* shape a polymicrobial community by acting as vehicles of public transport and that patterns within a microbial community arise as a result of this active motion. Over longer time scales, a combination of active motion and cell growth can shape the spatial features of a microbial community.

## Results & Discussion

### T9SS and its associated rotary motor are functional in gliding bacteria of the human microbiome

We found that many *C. gingivalis* cells were able to glide over a glass surface, moving at speeds of about 1 μm/s (Fig. 1A and Movie S1). Also, cells of *C. gingivalis* could tether to a glass surface and rotate about fixed axes at speeds of about 1.5 Hz (Fig. 1B, C and Movie S2). This behavior is similar to that of the environmental gliding bacterium *Flavobacterium johnsoniae*, where rotation of a cell around a fixed axis is attributed to the presence of a rotary motor associated with the T9SS that powers gliding (31). *sprA, sprE, sprT*, and *gldK*-*N* genes encode the proteins that form the core T9SS. *gldA*-B, *gldD*, and *gldF-J* encode proteins that associate with the T9SS to enable gliding motility. The bacterium *F. johnsoniae* has all 15 genes. T9SS is found in about 62% of the sequenced species of the phyla Bacteroidetes (32). *Capnocytophaga* species found in the human oral microbiome have all T9SS and gliding motility genes. Of the seven abundant non-motile bacteria used in this study, *P. endodontalis* and *P. oris* have T9SS genes but they lack the gliding motility genes (Fig. 1D). These observations combined with the presence of all T9SS-gliding motility genes and mobile cell-surface adhesins demonstrate that the machinery that enables gliding motility of members of the Bacteroidetes phyla is functional in *C. gingivalis*.

**Figure 1:**
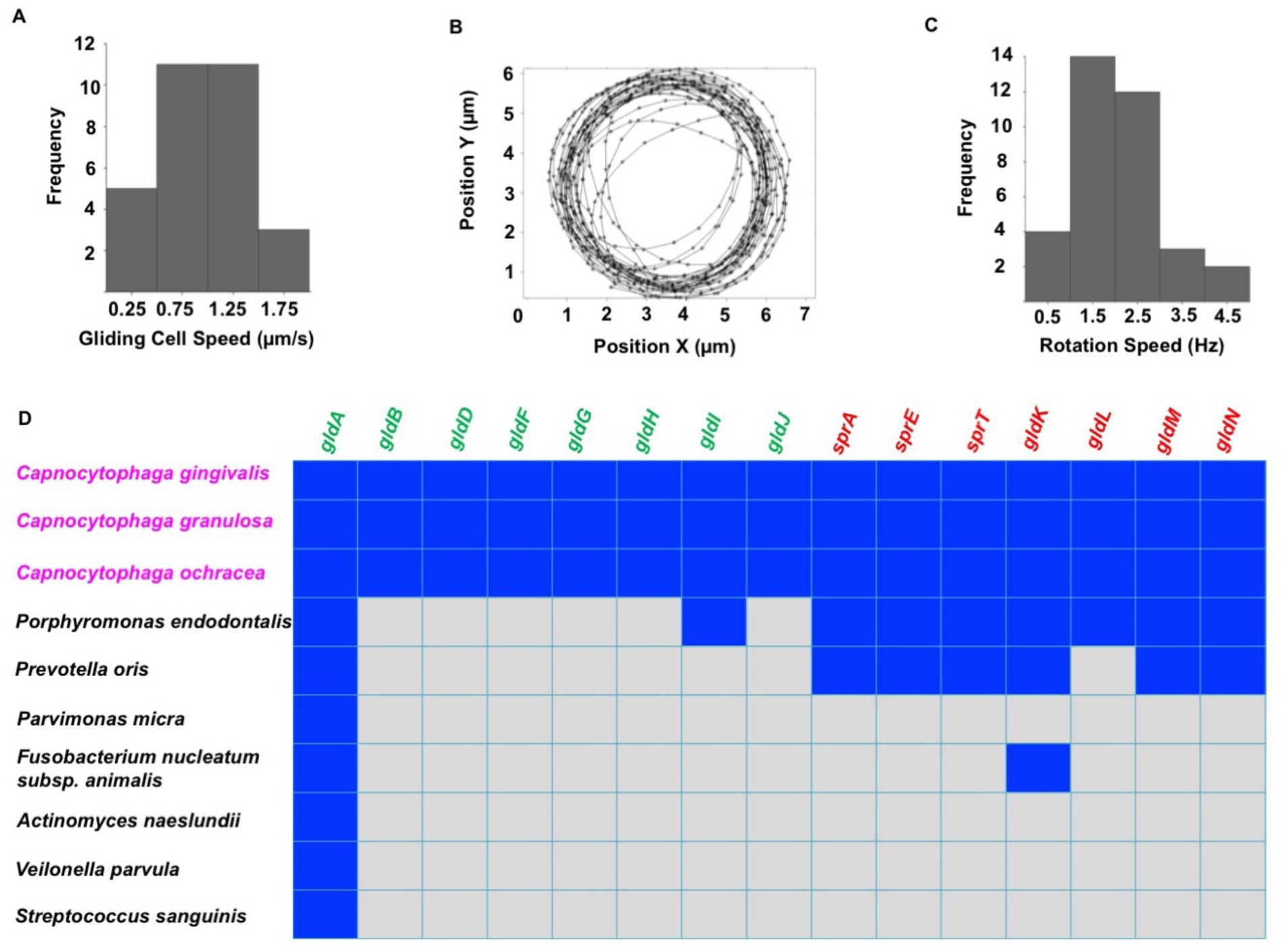
The type IX secretion system (T9SS) and its associated rotary motor are functional in gliding bacteria of the human microbiome. (A) The frequency distribution of gliding speeds of a population of *C. gingivalis* cells peaked around 1 μm/s. **(B)** The center of mass of a *C. gingivalis* cell tethered to glass followed a circular trajectory. **(C)** The frequency distribution of rotation speed of a population of tethered *C. gingivalis* cells peaked around 1.5 Hz. Experiments **A**-**C** done at room temperature on glass. **(D)** A matrix showing the similarity between genes that encode the T9SS and gliding motor proteins of abundant bacteria of the human oral microbiome. The reference gene source was *F. johnsoniae.* A blue square indicates presence while a grey square indicates the absence of a gene. BLAST was performed using the Human Oral Microbial Database (26). In order to eliminate random matches, an e value of e^−5^ was used as the threshold. The core gliding and core T9SS genes are shown in green and red, respectively. Bacteria shown in magenta have all green and red genes and can glide.

### Gliding bacteria from the human microbiome move collectively

We refer to the collective motion of gliding bacteria as “swarming” and to large groups of moving cells as “swarms”. Compact subgroups that move as a unit are called “slugs”. *C. gingivalis* cells displayed swarm behavior over the surface of agar. These swarms moved in a synchronized manner, and circular motion of cells was observed near the edge of the swarm. Their motion was tracked with micron-sized surfactant-encased gas bubbles. Such bubbles floated on the top layer of the swarm. (Fig. 2A-B and Movie S3). The gas bubbles near the leading edge of a swarm moved in circular trajectories predominantly in the counter-clockwise direction (Fig. 2C). The speed of bubbles, which is also the speed of the top layer of the swarm, was around 18 μm/min. (Fig. 2D).

**Figure 2.**
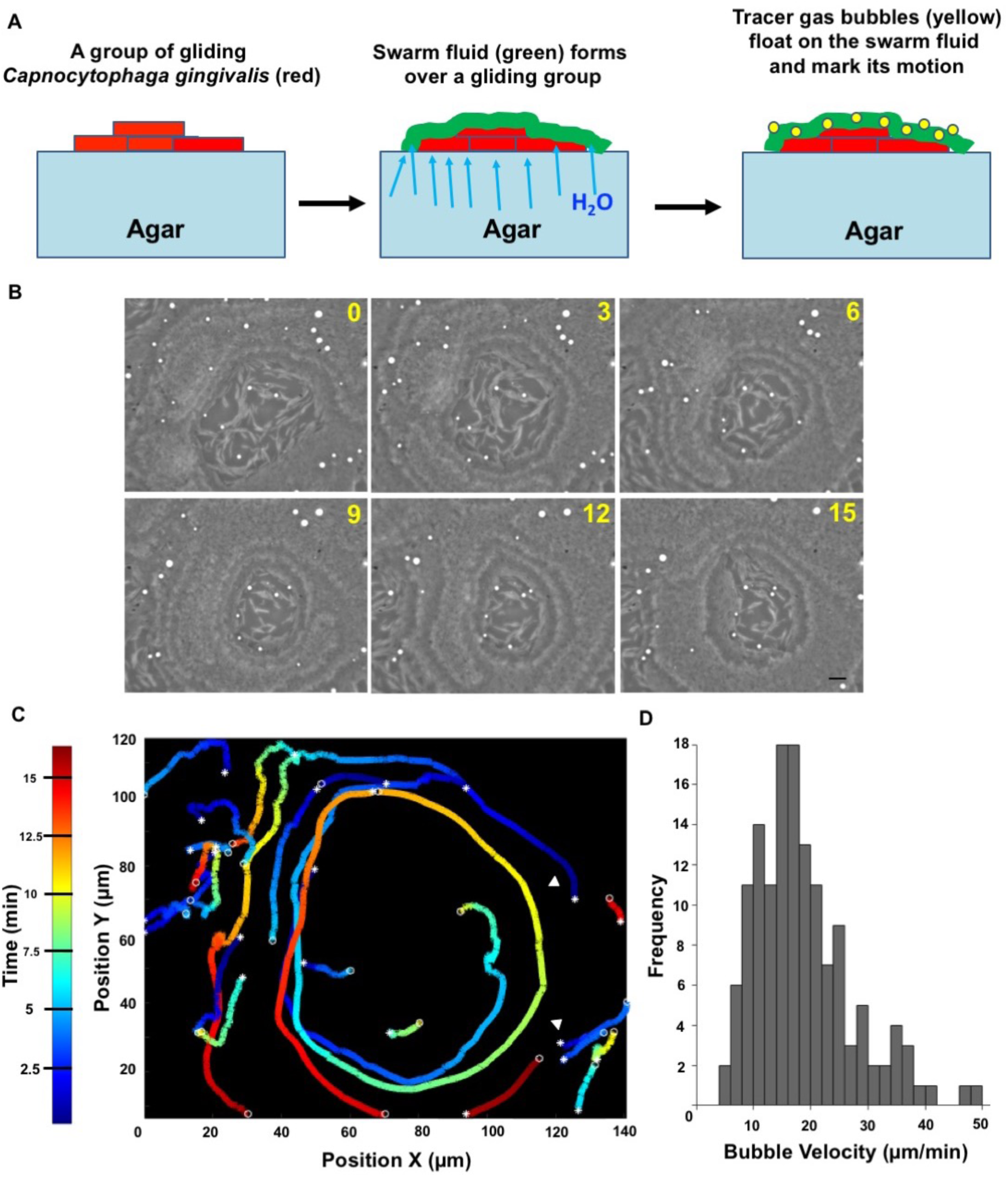
Synchronized swarm behavior of bacteria from the human microbiome. (A) A graphical representation of the use of gas bubbles to trace fluid flow patterns of a swarm. **(B)** Time-lapse images of a swarm with gas bubbles (white) moving on top. The time lapse is in minutes. Scale bar represents 10 μm. **(C)** Trajectories of the gas bubbles moving in a circular pattern on a swarm. Color map depicts time. **(D)** A frequency distribution of the speed of gas bubbles. Experiments **B**-**D** done at 37°C on an agar pad.

### Speeds of layers of a swarm are additive

The swarms of *C. gingivalis* formed layers which were superimposed on top of one other. If layers move in similar directions and cells in the bottom layer move with a speed *v*, then cells in the second layer would move with a speed *2v*. To test this hypothesis, a small percentage of fluorescently labeled *C. gingivalis* cells were mixed with unlabeled *C. gingivalis* cells. Fluorescently labeled cells were tracked in a swarm containing both labeled and unlabeled cells (Fig. 3 inset and Movie S4). The speed distribution of cells was bimodal with peaks around both 9 and 18 μm/min (Fig. 3A). Slugs that appeared near the edge of a swarm were tracked and their speed was around 9 μm/min (Fig. 3B-C and Movie S5). This suggests that the speed of a swarming cell and its synchronization correlates with its location in a three-dimensional spatial domain.

**Figure 3.**
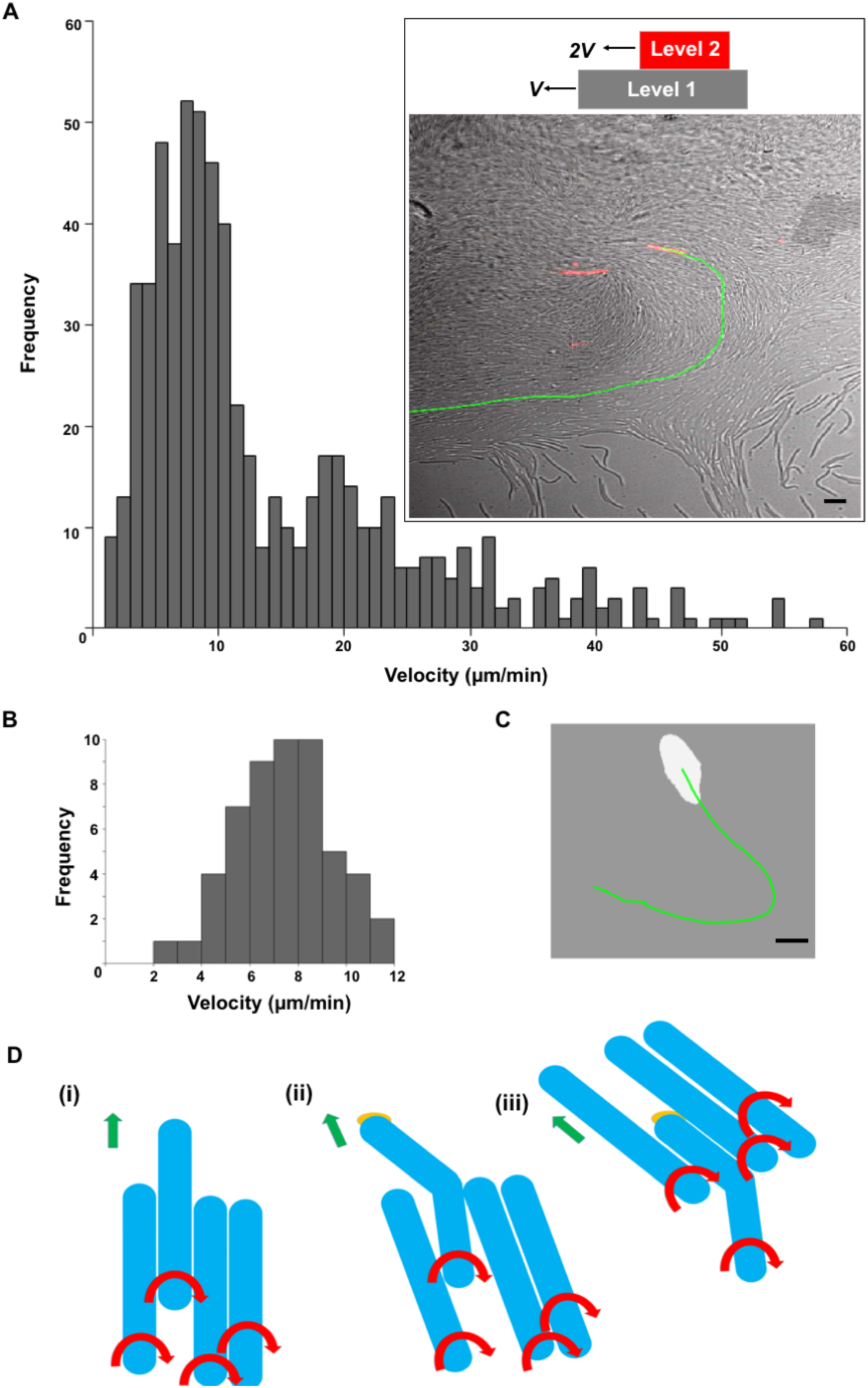
Speeds of layers of a swarm are additive. (A) A bimodal frequency distribution of the speed of fluorescently-labeled cells moving within a swarm. The inset shows a picture of a swarm with a fluorescently-labeled cell (red) and the trajectory of that cell (green). Scale bar represents 5 μm. **(B)** The frequency distribution of speed of slugs (a small group of mobile cells). **(C)** A picture showing a slug (white) with the trajectory of its motion in green. Scale bar represents 5 μm. Experiments A-C done at 37°C on an agar pad. **(D)** An explanation for how a group of cells moves counter-clockwise. (i) A group of flexible cells (blue) moving as right handed screws. Red arrows show the direction of rotation of the cell body and green arrows show the direction of motion of a group. (ii) The leading end of one cell sticks to an external surface via multiple adhesins (orange) while the rest of the cell body keeps twisting. This produces a torsion that bends the cell towards the left of the direction of motion of the group. (iii) Cells in a group orient themselves along the cell body of the bent cell and the group moves counter-clockwise.

### Counter-clockwise motion of a swarm can be explained from the right-handed motion of single cells

Single gliding cells move in a manner similar to a right handed screw. If the front end of a flexible gliding cell moving as a right handed screw sticks to an external surface, the cell body bends resulting in the front end of the cell pointing towards the left (Fig 3D and S2). Models that explain group behavior suggest that in a group, the direction of motion of an individual cell depends on the direction of motion of its neighbors and perturbations in the noise of the system result in varying patterns of a swarm (14). Cells that are in the vicinity of a bent cell move along the direction of the bend, and this direction of motion gets propagated in a swarm, thus resulting in its counter-clockwise motion (Fig. 3D).

### A cargo-transporter relationship exists between non-motile and gliding bacteria of the human microbiome

Genes that encode the bacterial T9SS and a cell surface adhesin SprB are present in *Capnocytophaga gingivalis*, which is abundant in human oral biofilms. *C. gingivalis* belongs to the phylum Bacteroidetes and is related to the model gliding bacterium *F. johnsoniae*. Addition of an antibody raised against the SprB of *F. johnsoniae* to a culture of *C. gingivalis* caused cell aggregation (Fig. S1), suggesting that *C. gingivalis* carries SprB on its surface. SprB is known to bind to polysaccharides, which are abundant on most cell walls. To test the hypothesis that this adhesin might bind to the cell-surfaces of non-motile bacteria, seven abundant and non-motile bacterial species found in human subgingival biofilms were co-cultured with gliding *C. gingivalis*. The non-motile bacteria were *Porphyromonas endodontalis, Prevotella oris, Parvimonas micra, Actinomyces sp.* oral taxon-*169, Fusobacterium nucleatum, Streptococcus sanguinis*, and *Veilonella parvula*. Single cells of these non-motile species attached to the cell-surface of *C. gingivalis* and were transported as cargo along a *C. gingivalis* cell from one pole to the other (Fig. 4A and Movie S6-S12). The adhesin SprB of *F. johnsoniae* moves in a spiral fashion on the cell-surface (17). A non-motile bacterium on the surface of a *C. gingivalis* cell moves in a similar way (Fig. 4B).

**Figure 4.**
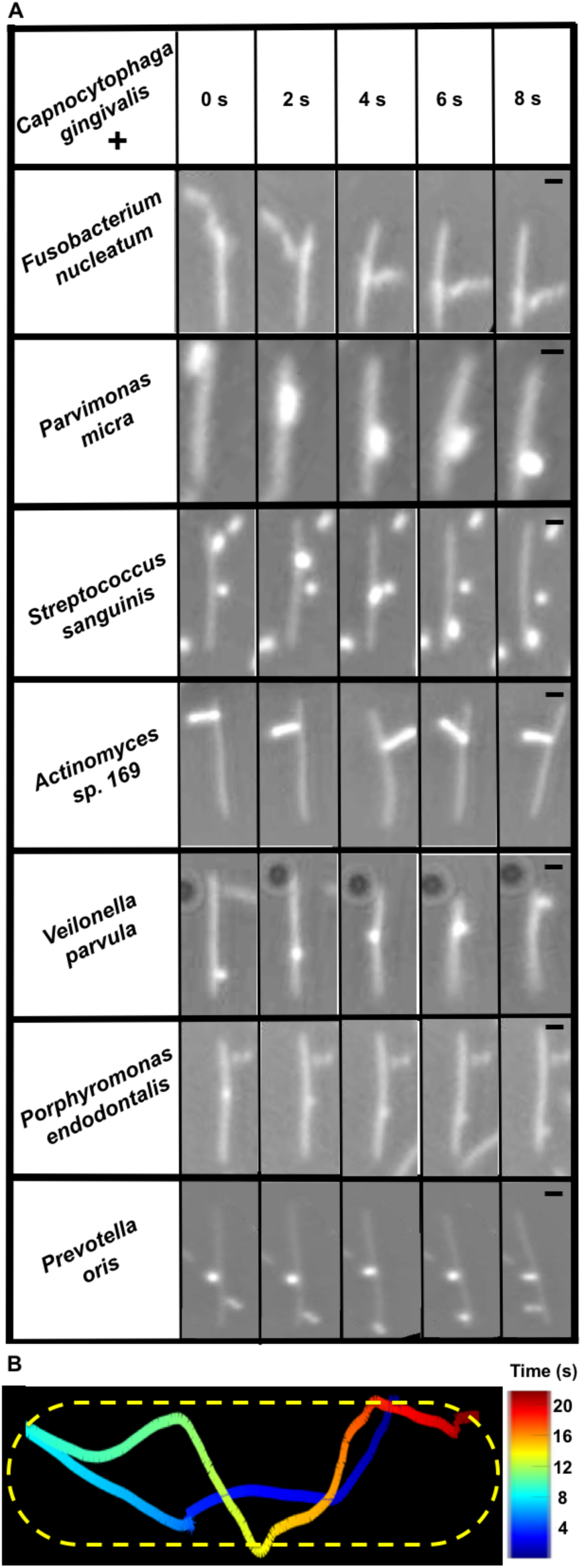
Cargo-transporter relationship exists between non-motile and gliding bacteria of the human microbiome. (A) Seven different species of non-motile bacteria were carried along the surface of *C. gingivalis* cells, moving from one end of the cell to the other. The images have been aligned to reveal the motion of the cargo relative to the *C. gingivalis* cell. The raw data are shown in Movies S6-S12. Scale bar represents 1 µm. The eight bacteria shown in the matrix are some of the abundant microbial species found in a human subgingival biofilm. **(B)** The trajectory of a non-motile bacterium *Prevotella oris* transported along the length of a *C. gingivalis* cell shown as a detailed example of the cargo-transporter relationship described above. *P. oris* moved from one pole to the other and looped back after it reached a pole. The *C. gingivalis* cell is outlined by a dotted line; color map indicates time. Experiments done at room temperature on glass.

### Long-range cargo transport shapes a microbial community

The spatial localization of bacteria was studied in a polymicrobial community comprising *C. gingivalis* and the seven non-motile bacteria used for the experiments of Fig. 4A. To explore if *C. gingivalis* swarms had the capability to transport the non-motile bacteria over long distances, the cell surfaces of the non-motile bacteria were labeled using seven different amino-specific succinimidyl ester Alexa Flour fluorescent dyes. Physiological features, such as the motility of labeled *C. gingivalis* remained unaltered, thus providing a powerful tool for live-cell imaging. A polymicrobial community containing seven fluorescently labeled non-motile bacteria and gliding *C. gingivalis* was imaged, and spectral imaging was used to separate the fluorescent signals. Long-range transport of non-motile bacteria by swarming *C. gingivalis* was observed in a polymicrobial community (Fig. 5A and Movie S13). Within 20 min, the polymicrobial community evolved in such a way that many non-motile bacteria were arranged as islands surrounded by swarms of *C. gingivalis* and some non-motile bacteria (Fig. 5A). A control, in which *C. gingivalis* was de-energized by prior treatment with sodium azide exhibited a uniform distribution (Fig. 5B). The non-motile bacteria were actively transported via the swarms as cargo (Fig. 5C). Polymicrobial aggregates of no-nmotile bacteria that were transported by *C. gingivalis* changed their structures while moving with a swarm (Fig. 5D and Movie S14). Tracking of individual bacterial species revealed that some non-motile bacteria moved distances of up to 30 µm in a swarm. Some non-motile bacteria, such as *Fusobacterium nucleatum*, used this mode of transport more efficiently than others (Fig. 6A–E). Since equal numbers of cells of each species were included in the mixture added to the agar pad (see Methods) the transport efficiency, the relative number of tracks for each species that appear in Fig. 5C, can be found from the areas of the distributions in Fig. 6. Additional information can be obtained from the lengths of distribution tails. Differences in the composition of the polysaccharides or cell-surface proteins of the non-motile bacteria could have an impact on their ability to be transported as cargo. Based on our results, we propose a model where a microbial community containing gliding bacteria can use surface motility and cargo transport to find a specific niche. Once a niche is established, the bacteria can arrange in well-defined spatial structures (Fig. 7). Evidently, cargo transport plays a role in forming the spatial features of polymicrobial biofilms that contain *Capnocytophaga sp*. Such biofilms are widespread in the human gingival regions and the tongue. It is known that cooperation in the form of mutualism amongst bacterial species occurs via the sharing of public goods (33). We demonstrate that public transport enables bacterial cooperation and shapes communities of bacteria found in the human body.

**Figure 5.**
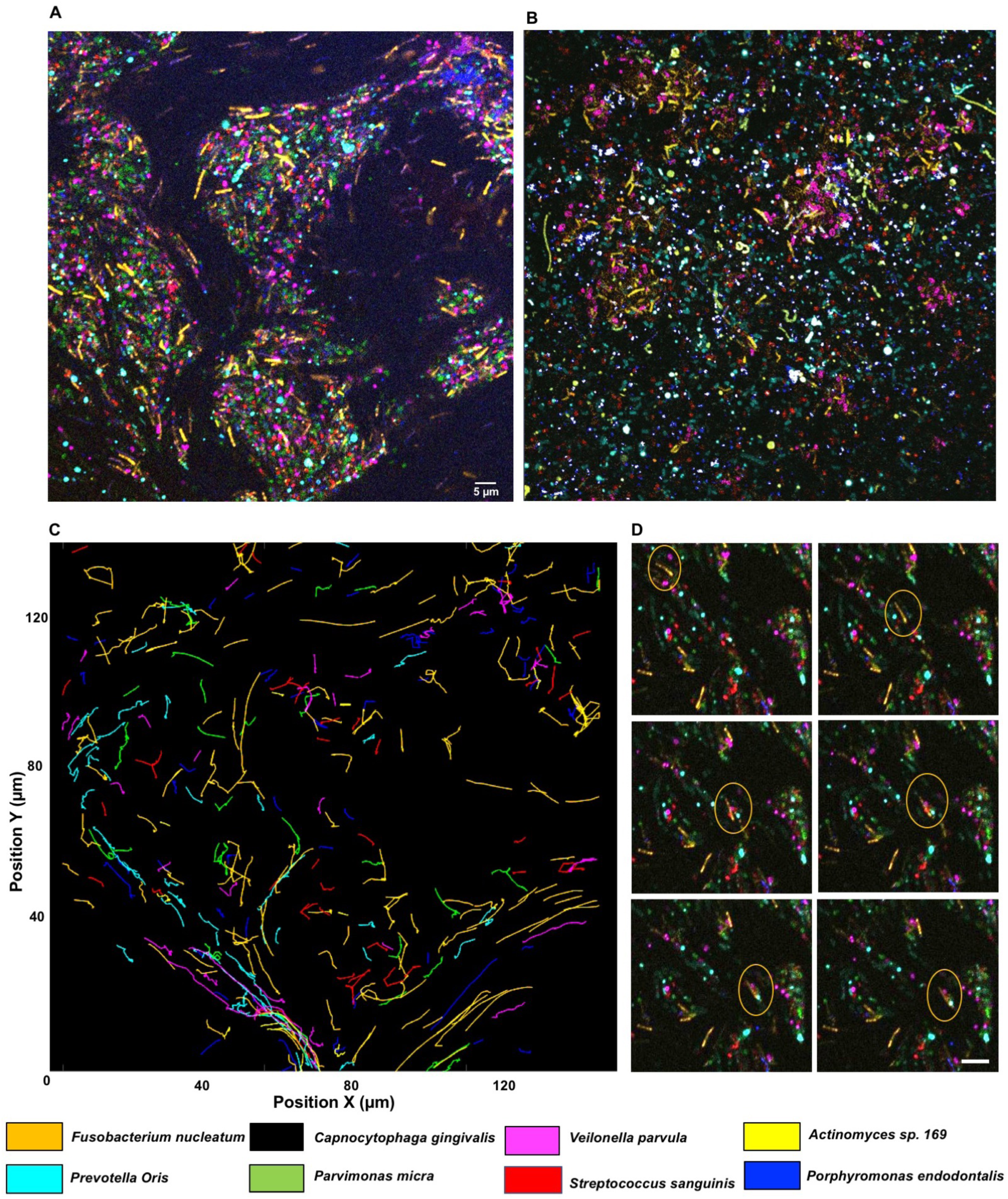
Long-range cargo transport shapes a microbial community. (A) A microbial community containing fluorescently-labeled bacteria. Eight bacteria (seven non-motile and one motile) of the human microbiome attained a shape in which the non-motile bacteria were organized in islands by the swarming bacteria. This is a still image from the movie S13. **(B)** A control in which the experiment of panel A was repeated with de-energized cells of *C. gingivalis*. Scale bar represents 5 µm. Experiments done at 37C on an agar pad. **(C)** 20-minute trajectories of the displacement of non-motile bacteria showing that the swarming microbe transports non-motile bacteria over long distances. **(D)** Time-lapse images taken at an interval of 1 minute of a small region from Movie S13 showing that dynamically forming polymicrobial aggregates of non-motile bacteria move long distances in a swarm.

**Figure 6. (A-G).**
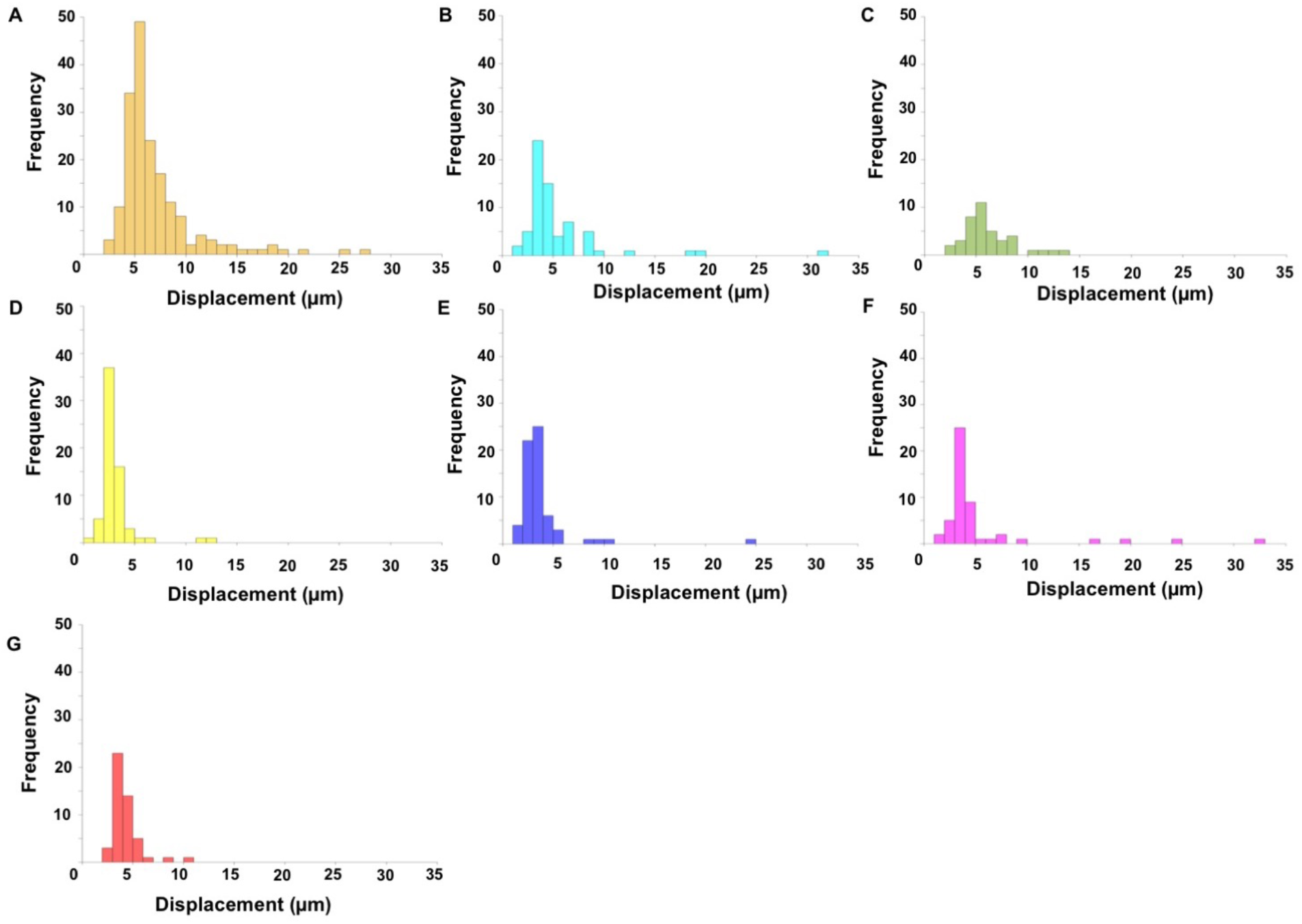
Frequency distributions of the distances that 7 species of non-motile bacteria were transported by a swarm of *C. gingivalis*. Different species are indicated in the color code defined in Fig. 4. The frequency distributions are for cells whose tracks appear in Fig. 5B.

**Figure 7.**
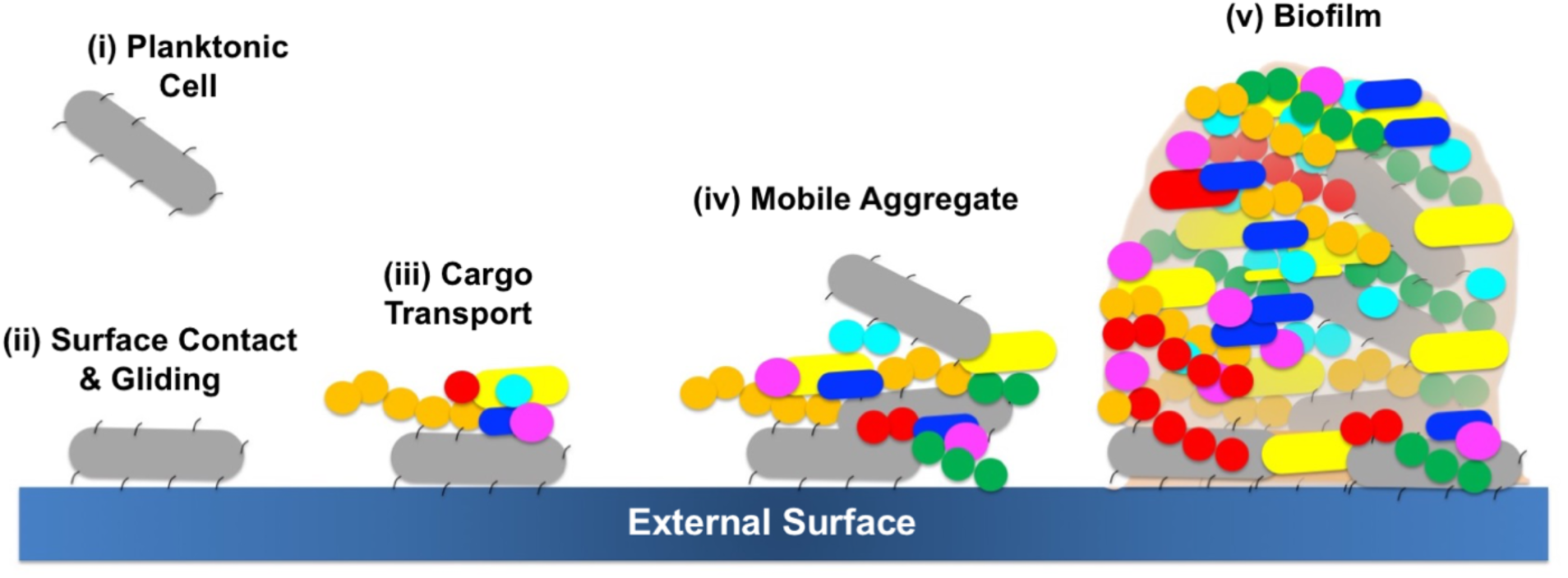
A model depicting how a gliding bacterium from the human microbiome can carry polymicrobial cargo, form mobile aggregates, and find a spatial niche where a polymicrobial biofilm can develop.

## Methods

### Strains and growth

Cells of *Capnocytophaga gingivalis* ATCC 33624 were used for gliding motility assays. The strain was originally isolated from the periodontal lesion of a human subject. *C. gingivalis* cells were motile when grown on TSY agar, which contains trypticase soy broth (TSB) 30 g/L, yeast extract 3 g/L, and 1.5% Bacto Agar (Difco). Growth was carried out in a CO_2_-rich anaerobic environment at high relative humidity. This was achieved by placing the inoculated agar plates in an AnaeroPack System jar (Mitsubishi Gas Chemical Co.) containing 3 candles and an ignited sheet of Kimtech paper at 37˚C for 48 h. Non-motile bacteria were originally isolated from the oral cavity of human subjects and they are cataloged in the Human Oral Microbiome Database (26). *Fusobacterium nucleatum, Actinomyces sp. oral taxon-169, Parvimonas micra, Streptococcus sanguinis, Prevotella oris*, and *Veilonella parvula* were grown on a medium containing tryptic soy broth 20 g/L, brain heart infusion broth 26 g/L, yeast extract 10 g/L, hemin 5 mg/L, 5% sheep blood, and agar 22 g/L at 37˚C in an anaerobic chamber. *Porphyromonas endodontalis* was grown on a medium containing TSB 37 g/L, hemin 5 mg/L, NaHCO_3_ 1 g/L, yeast extract 1 g/L, 5% sheep blood, and 1.5% agar at 37˚C in an anaerobic chamber.

### Single cell motility assay

*C. gingivalis* cells were suspended in TSY broth which contains trypticase soy broth (TSB) 30 g/L and yeast extract 3 g/L. The cell suspension was injected into a tunnel slide (the space between a slide and a coverslip separated by one layer of double-stick Scotch tape) and was allowed to stand for 5 min. Then, cells in the tunnel slide were washed with 0.25% methyl-cellulose (Methocel 90 HG Sigma-Aldrich catalog number 64680; viscosity of a 2% solution in water at 20˚C about 4000 cP) containing TSY broth. After another 5 min, the gliding cells were imaged using a Nikon Plan 40x BM na 0.65 objective (Nikon, Melville, NY) and a ThorLabs DCC1545 M-GL camera (ThorLabs, Newton, NJ). Images were analyzed using a custom MATLAB (The MathWorks, Natick, MA) code. To image and analyze tethered and rotating cells, a suspension of *C. gingivalis* cells was injected into a tunnel slide and washed with TSY broth. Some of the cells tethered to glass and displayed rotation. Images of such cells were captured and analyzed using the method described above.

### Generation of microbubbles and tracking fluid flow

The method for producing surfactant-encapsulated air bubbles is given in (34). About 70 μL water-insoluble surfactant Span 83 (Sorbitan sesquioleate, S3386; Sigma-Aldrich) was mixed with 100 mL sterile deionized water in a glass bottle with two inlets. Air was injected into the mixture for 1 h, turning the solution into a milky white suspension. To prepare an agar pad, about 700 µl of melted 1% TSY agar was applied to a glass slide (#48300–026, VWR, 25×75 mm, 1.0 mm thick). A 1 μL sample of the Span 83 suspension was placed on top of the agar pad and the slide was placed in a high relative-humidity chamber at 37°C for about 40 min. After the water in the sample was absorbed by the agar, the Span 83 droplet spread out explosively and soon contracted into an array of micron-sized air bubbles with surfactant walls. Then, 1 μL of *C. gingivalis* cells suspended in sterile deionized water were added near the edge of the area containing the surfactant. After incubation for about 20 min, *C. gingivalis* started to swarm and reached the area with the bubbles. The microbubbles formed by the surfactant were used as tracers to image the patterns of motion of the top layer of a swarm. Microbubbles were imaged using the Nikon microscope and ThorLabs camera system described above and their motion was tracked using ImageJ and a custom MATLAB script.

### Cargo transport and polymicrobial interactions

In order to study cargo transport by single cells, aliquots of cultures of *C. gingivalis* at O.D. 0.6 were mixed with similar aliquots of cultures of each non-motile species. The mixture of cell pairs was incubated for 10 min after which they were added to a tunnel slide, incubated for 5 min, washed with 0.25 % methyl-cellulose (defined above) containing TSY broth, and imaged using the Nikon microscope and ThorLabs camera system described above. Motion of cargo cells on the transporter cells was tracked using ImageJ and the motion of the transporter cell was tracked using a custom MATLAB script. To obtain the trajectory of a moving cargo cell with respect to the transporter cell, the motion of the transporter cell was subtracted from the motion of the cargo cell using a custom MATLAB script. To test if the adhesin SprB is present on the surface of a *C. gingivalis* cell, 5 μL of affinity-purified antibody raised against *F. johnsonaie* SprB was added to 100 μL of *C. gingivalis* cells suspended in phosphate-buffered saline (PBS). After incubation for 10 min at 25˚C, the preparation and control with 5 μL PBS were injected into tunnel slides and were imaged using the Nikon microscope and ThorLabs camera system described above.

To image collective motion in a polymicrobial community, the cell-surface of live bacterial cells was stained with an Alexa Fluor NHS ester (succinimidyl ester) dye (35). The cell suspensions were at O.D. 0.6. The following bacteria-dye combinations were used: *Porphyromonas endodontalis*: Alexa 514, *Prevotella oris*: Alexa 555, *Parvimonas micra*: Alexa 568, *Actinomyces sp. oral taxon-169*: Alexa 594, *Fusobacterium nucleatum*: Alexa 633, *Streptococcus sanguinis*: Alexa 647, *Veilonella parvula*: Alexa 660. *C. gingivalis* was not labeled. Labeling was performed by mixing 1 μL of 4 μg/μL dye with 10 μL of the respective bacterial cell suspension, incubated in the dark and shaking gently for 1 h and washed twice with sterile deionized water by centrifugation to remove cell-free dye. Equal volumes of cultures at the same O.D. (of *C. gingivalis* and the seven fluorescently-labeled bacteria) were mixed and 1 μL of the mixture was added to an agar pad and incubated for 20 min at high relative humidity. To maintain an anaerobic environment, the agar pad was sealed with a cover glass and paraffin wax. After the incubation, swarming was observed. Cells were imaged with a Zeiss LSM 880 Airyscan Confocal microscope in a chamber set at 37˚C. Bacteria labeled with the dyes described above were used for calibration, and spectral imaging was used to separate the 7 fluorescent signals. The images were analyzed using ImageJ and superimposition of the fluorescent signal onto a differential interference contrast (DIC) image provided the location of *C. gingivalis*. The motion of individual bacteria was tracked using ImageJ and was analyzed using a custom MATLAB script. The above experiment was repeated using cells of *C. gingivalis* that had been de-energized by treatment with 1 mM sodium azide for 10 min followed by 2 washes with deionized water.

### Imaging of fluorescent cells in a swarm

1 μL of *C. gingivalis* cells at O.D. 0.6 were added to an agar pad and incubated at 37 °C for 20 min in a chamber at high relative-humidity. The edge of a swarm had smaller motile groups of cells (slugs) which were imaged using the Nikon microscope and ThorLabs camera system described above. Motion of the slugs was tracked using a custom MATLAB script. An aliquot of *C. gingivalis* cells suspended in sterile deionized water was labeled with Alexa Fluor 633 as described above. The labeled cells were mixed with unlabeled cells so that only 1% of the cells in the final mixture were labeled. This mixture was placed on an agar pad and fluorescently-labeled cells were imaged using a Zeiss LSM 880 Airyscan Confocal microscope. Motion of the cells was tracked using a custom MATLAB script.

## Acknowledgements

This research was supported by an NIH-NIDCR K99/R00 Pathway to Independence Award DE026826 to A.S. and an NIH-NIAID R01 AI016478 grant to H.C.B. We thank the Harvard Center for Biological Imaging for infrastructure and support. We thank Karen A. Fahrner for comments on the manuscript.

---

**Figure S1:**
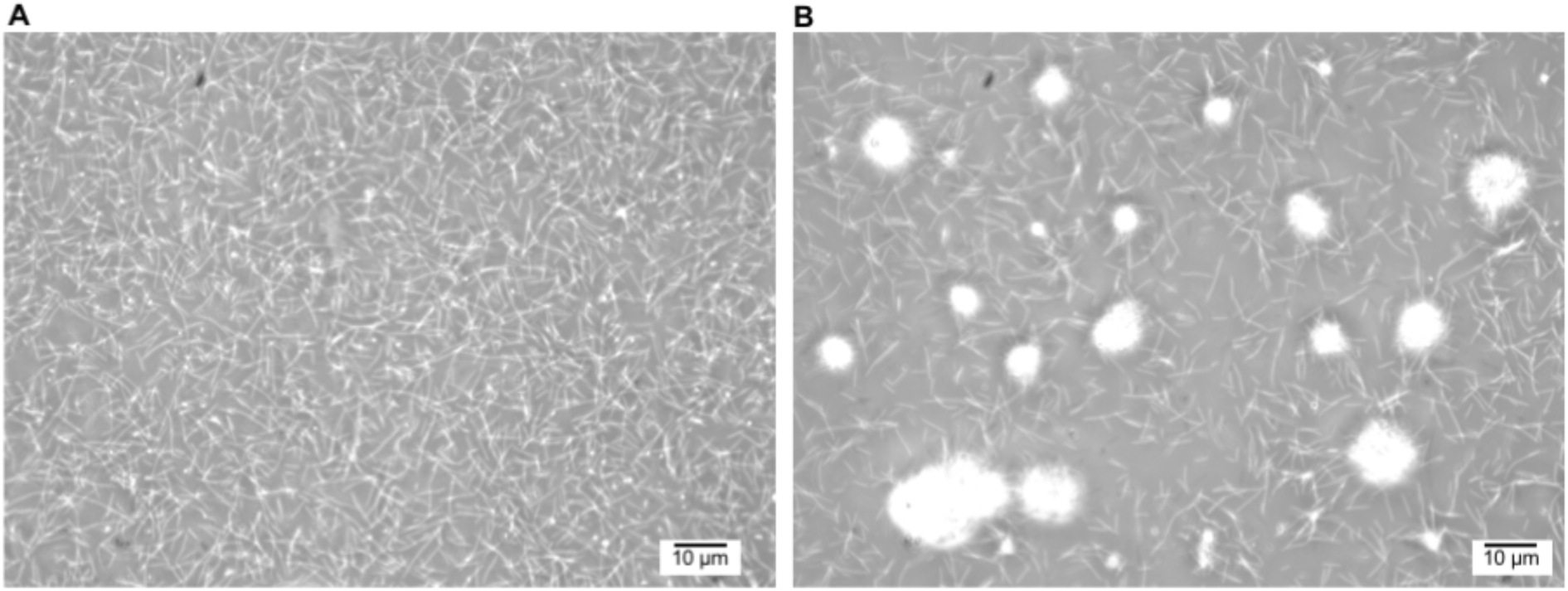
*F. johnsoniae* SprB antibody causes aggregation of *C. gingivalis* cells. **(A)** *C. gingivalis* cells with no antibody **(B)** *C. gingivalis* cells with anti-SprB antibody. Experiments done at room temperature.

**Figure S2:**
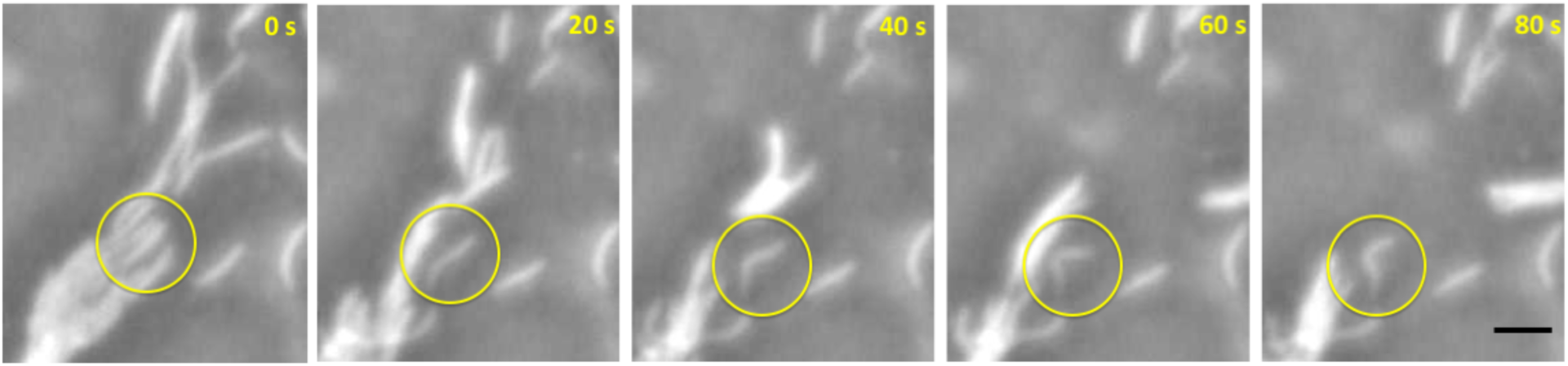
Images showing one stuck and bent cell (marked by a yellow circle) within a small group (slug) of *C. gingivalis*. Experiments done at room temperature. Scale bar represents 5 μm.

## Supplementary Movies

**Movie S1.** Gliding of single *C. gingivalis* cells over a glass surface.

**Movie S2.** *C. gingivalis* cell tethered to a glass surface and rotating about a fixed axis.

**Movie S3.** A swarm of *C. gingivalis* moving in a circular fashion with layers on top of one another. Gas bubbles (bright white spheres) move along the top layer of the swarm. A less dense region with small mobile aggregates is seen near the center of the swarm.

**Movie S4.** Fluorescently-labeled *C. gingivalis* cells in a swarm containing both labeled and unlabeled *C. gingivalis* cells.

**Movie S5.** Motion of slugs found near the edge of a swarm.

**Movie S6.** *P. oris* transported as cargo along the length of a single *C. gingivalis* cell.

**Movie S7.** *F. nucleatum* transported as cargo along the length of a single *C. gingivalis* cell.

**Movie S8.** *P. micra* transported as cargo along the length of a single *C. gingivalis* cell.

**Movie S9.** *S. sanguinis* transported as cargo along the length of a single *C. gingivalis* cell.

**Movie S10.** *Actinomyces sp. 169* transported as cargo along the length of a single *C. gingivalis* cell.

**Movie S11.** *V. parvula* transported as cargo along the length of a single *C. gingivalis* cell.

**Movie S12.** *P. endodontalis* transported as cargo along the length of a single *C. gingivalis* cell.

**Movie S13.** A polymicrobial community with seven non-motile fluorescent bacteria being moved by a *C. gingivalis* (grey) resolved via spectral imaging. See Fig. 5A for a color key.

**Movie S14.** Time-lapse images of a small region from Movie S13 shows that a polymicrobial aggregate of non-motile bacteria moves long distances in a swarm. For better contrast, *C. gingivalis* cells are represented in black while non-motile bacteria are represented by the color key shown in Fig. 5.

